# *Brucella* proline racemase protein A targets Tpl2 to promote IL-10 secretion for establishment of chronic infection

**DOI:** 10.1101/2025.02.10.637385

**Authors:** Huan Zhang, Yuanzhi Wang, Yueli Wang, Huilin Hou, Yaqian Wang, Xiaoyu Deng, Zhenyu Xu, Xujin Xia, Mingguo Xu, Zhen Wang, Yong Wang, Changsuo Zhang, Kait Zhumanov, Jinliang Sheng, Hui Zhang, Zhongchen Ma, Jihai Yi, Chuangfu Chen

**Affiliations:** School of Animal Science and Technology, Shihezi University, 832000-Shihezi City, Xinjiang, China; School of Medicine, Shihezi University, 832000-Shihezi City, Xinjiang, China; Center for Animal Disease Control and Diagnosis in Bayinggolin Mongol Autonomus Prefecture, 841000-Korla City, Xinjiang, China; Tecon Biological Co. Ltd., 830011-Urumqi, China; School of Animal Science and Technology, Kazakh National Agrarian University, 040900-Karasai raion, Almaty, Kazakhstan

**Keywords:** *Brucella melitensis* M5-90, proline racemase, tumor progression locus 2, IL-10, vaccine

## Abstract

IL-10, an anti-inflammatory cytokine, plays a crucial role in limiting immune responses to pathogens, preventing host damage. However, the mechanisms underlying *Brucella*-mediated IL- 10 production remain incompletely understood. In this study, we demonstrate that the proline racemase protein A (PrpA) of *Brucella melitensis* M5-90 induces macrophages to secrete IL-10 by activating the Tpl2-ERK signaling pathway, thereby promoting chronic infection. Moreover, *Tpl2* deletion impairs macrophage bactericidal ability, accompanied by reduced TNF-α and IL-1β but unaffected IL-10 levels. Additionally, Trp309, Glu103, and Glu129 of PrpA participate in interaction with Tpl2, but these residues do not influence PrpA-mediated IL-10 production in macrophages. *PrpA* deletion enhances IFN-γ levels, specific anti-*Brucella* IgG, and CD4+ and CD8+ T cell numbers in mice. Furthermore, the *Brucella melitensis* M5-90 *prpA* mutant provides higher protection than the parental strain against virulent *Brucella melitensis* M28 infection in mice. Our findings suggest that *Brucella* PrpA promotes IL-10 secretion by macrophages through Tpl2 activation for bacterial survival and persistent infection, making the *Brucella melitensis* M5-90 *prpA* mutant a promising vaccine for enhanced protection.

**Author Summary:** IL-10, an anti-inflammatory cytokine, is exploited by *Brucella* for survival. We identified *Brucella* PrpA as a potent IL-10 inducer, activating the Tpl2-ERK signaling pathway. *Tpl2* deletion increased *Brucella* survival in macrophages, accompanied by reduced TNF-α and IL-1β but unaffected IL-10 secretion. Residues (Trp309, Glu103, and Glu129) of PrpA were crucial for interaction with Tpl2, yet did not impact PrpA-mediated IL-10 production. The *B. melitensis* M5-90 *prpA* mutant induced higher anti-*Brucella* IgG, IFN-γ, and CD4+ and CD8+ T cell numbers in mice, providing better protection than *B. melitensis* M5-90. These findings unveil PrpA’s role in IL-10 production and highlight the potential of the *B. melitensis* M5-90 *prpA* mutant as a promising vaccine for further evaluation.

## Introduction

Brucellosis, resulting from *Brucella spp*. infection, stands globally recognized as a significant zoonosis. With an annual incidence exceeding 500,000 new reported human cases worldwide, its substantial public health implications are evident(1). Human brucellosis manifests initially as an acute undulating fever, progressing into a chronic disease often accompanied by debilitating symptoms, including arthritis, endocarditis, meningitis, and spondylitis(2, 3). In livestock, brucellosis leads to abortion and infertility, causing substantial economic losses in husbandry, estimated at 16,386,500 yuan (2,589,067 USD) annually in China(4, 5). Regions heavily dependent on animal production for economic sustenance particularly feel the pronounced impact of brucellosis-induced losses.

Innate immune responses serve as the primary defense against bacterial pathogens. However, immune suppression during persistent bacterial infections poses significant public health challenges(6–8). *Brucella* employs various strategies to interfere with or evade immune recognition and suppress immune responses(9–11). Notably, *Brucella* inhibits innate immune signaling using proteins like Btp1 or TcpB containing TIR domains(12–14). Additionally, its resistance to complement activation and the production of a smooth, minimally endotoxic lipopolysaccharide (LPS) diminish its recognition by the Toll-like receptor 4, an innate immune sensor(15). Research suggests that *Brucella*’s Outer Membrane Protein 25 (OMP25) inhibits pro-inflammatory cytokine production by facilitating ubiquitination and degradation of Toll-like receptors (TLRs) and their adaptor proteins, contributing to compromised immune responses and persistent *Brucella* infection(16).

Interleukin-10 (IL-10) acts as an anti-inflammatory cytokine, crucial for preventing inflammatory and autoimmune pathologies. However, bacterial pathogens exploit IL-10 to suppress immune responses for their survival. *Salmonella*’s SarA effector protein induces IL-10 production in macrophages via STAT3 activation, supporting bacterial survival(17). Similarly, *Mycobacteria* Rv2145c triggers IL-10 secretion in macrophages through STAT3 activation, enhancing intracellular survival(7). Previous studies indicate that *Brucella* also utilizes IL-10 to promote intracellular survival and establish chronic infection(8, 18–21). Specifically, the proline racemase (PrpA) of *Brucella abortus* 2308 emerges as a potent IL-10 inducer. PrpA stimulation leads to IL- 10 secretion, and its removal diminishes *Brucella*’s ability to establish chronic infection in mice(20). Furthermore, PrpA, secreted into the macrophage cytoplasm, palmitoylated by the host cell, interacts with the NMM-IIA receptor, inducing IL-10 production in these macrophages(22–24). PrpA, therefore, stands as a crucial virulence factor enabling *Brucella* to modulate the immune response for chronic infection. Yet, potential additional mechanisms underlying PrpA-induced IL- 10 production in macrophages remain unknown.

The utilization of *Brucella melitensis* M5-90 (*B. melitensis* M5-90) in China for the prevention of brucellosis in sheep and goats is firmly established, providing effective protective immunity for these animals. However, the lingering virulence of *B. melitensis* M5-90 poses a potential risk of inducing abortion in gestational animals. Therefore, the identification and characterisation of virulence factors associated with *B. melitensis* M5-90 can serve as the foundation for developing safer and more efficacious vaccines(25). Previous studies have indicated that *B. abortus* 2308 PrpA stimulates splenocytes and macrophages to secrete IL-10, thereby modulating immune responses to establish chronic infection(20, 22–24). Consequently, PrpA may perform a similar function in *B. melitensis* M5-90 as it does in *B. abortus* 2308 during infection. The objectives of this study are as follows: 1) to explore the functions of *B. melitensis* M5-90 PrpA in macrophages; 2) to investigate other underlying mechanisms by which PrpA induces IL-10 production in macrophages; 3) to assess the immune response induced by the *B. melitensis* M5-90 *prpA* mutant in mice.

## Results

### *B. melitensis* M5-90 PrpA Protein Induces IL-10 In *Vitro* and In *Vivo*, alongside the capability to induce macrophage proliferation

Previous investigations have established that the PrpA protein of *B. abortus* 2308 has the ability to prompt murine naïve splenocytes and macrophages to secrete IL-10, contributing to the establishment of chronic infection(20, 22–24). To explore whether the PrpA protein in *B. melitensis* M5-90 retains the effect already described previously, and three other virulence factors of *Brucella*—wadC(26), OMP25(16), and RomA(27) were also selected for investigation. The recombinant forms of these proteins were expressed, purified, and confirmed through SDS-PAGE, revealing molecular weights of approximately 27 kDa for wadC, 30 kDa for RomA, 31 kDa for OMP25, and 30 kDa for PrpA (S1A Fig). Protein specificity was further validated using Western blotting (S1B Fig). Murine macrophages were individually stimulated with these four recombinant proteins, demonstrating that PrpA significantly induces IL-10 production in macrophages. Although the wadC can also slightly induce macrophages to secrete IL-10, the level of the IL-10 induced with the stimulation of wadC in macrophages was lower than that of the PrpA (Fig 1A). However, RomA and OMP25 did not elicit IL-10 secretion from macrophages. To comprehensively evaluate the IL- 10-inducing capacity of these proteins, Balb/c female mice were immunized with each protein. ELISA analysis of serum IL-10 levels revealed a significant enhancement in in *vivo* IL-10 production in response to PrpA. While OMP25 induced IL-10 production at 15 days post-immunization, the levels were lower than those induced by PrpA in *vivo* (Fig 1B). To rule out potential factors (e.g., residual LPS or low purity of the rPrpA protein) stimulating macrophages to secrete IL-10, a pcDNA3.1-*prpA* plasmid was constructed and transfected into macrophages, demonstrating that 3.5 μg or 4.5 μg of the plasmid significantly induced IL-10 production in macrophages (Fig 1C). Furthermore, PrpA did not induce macrophages to produce TNF-α (Fig 1D).

**Fig. 1:**
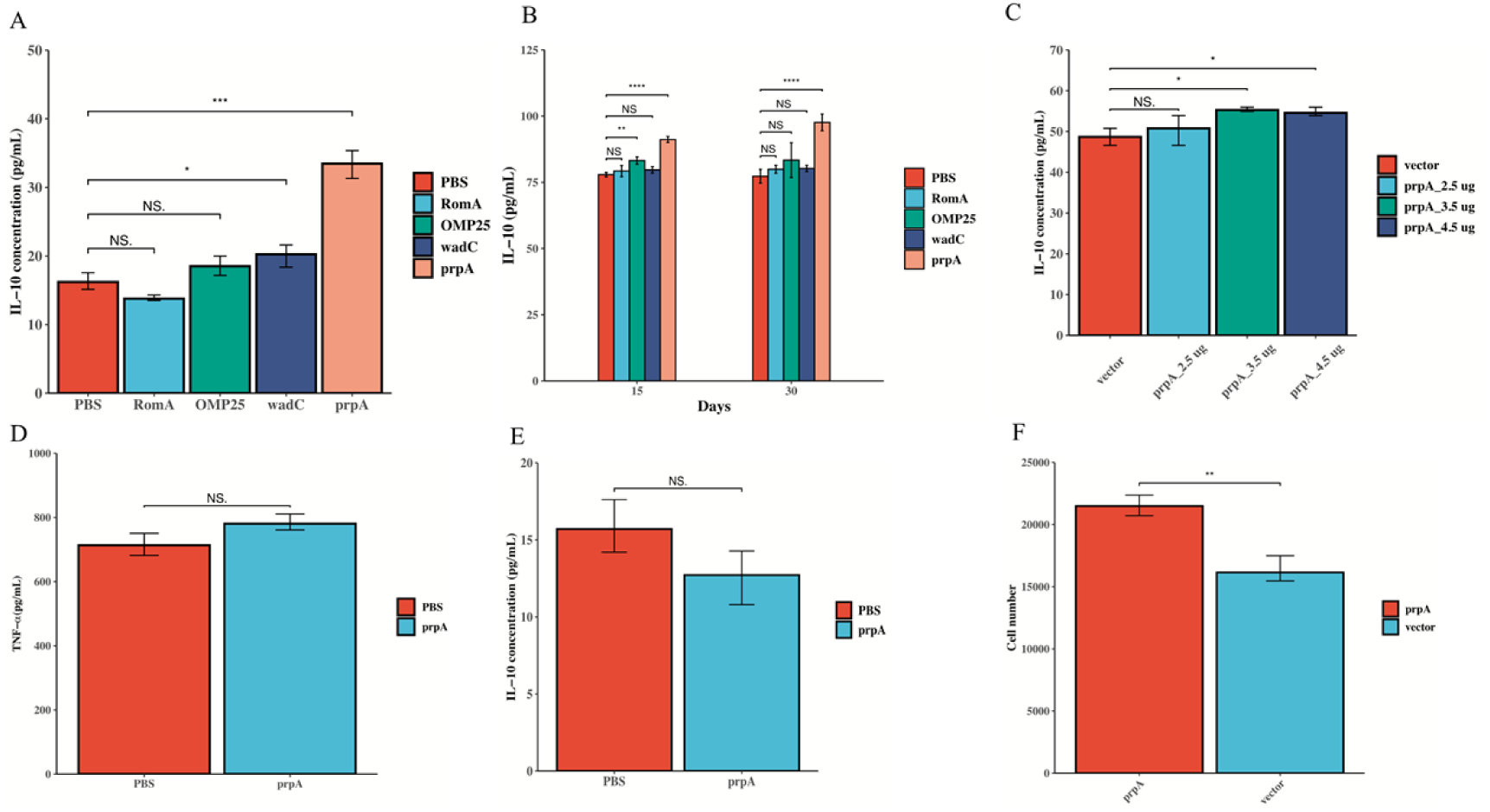
Investigation of RomA, OMP25, wadC, and PrpA in Inducing IL-10 Production. A) Evaluation of IL-10 concentration in macrophages post-stimulation with rRomA, rOMP25, rwadC, and rPrpA proteins in vitro. B) Measurement of IL-10 concentration in mice after immunization with rRomA, rOMP25, rwadC, and rPrpA proteins. C) Assessment of IL-10 concentration in macrophages post-transfection with varying doses of pcDNA3.1-*prpA* plasmid. D) Determination of TNF-α concentration in macrophages following stimulation with rPrpA protein. E) Measurement of IL-10 concentration in DC2.4 cells after stimulation with rPrpA protein. F) Illustration of PrpA’s role in promoting macrophage proliferation.

### Dendritic Cell Response to PrpA Protein and PrpA’s Role in Macrophage Proliferation

Dendritic cells (DCs) play a pivotal role in the innate immune response against *Brucella*. To explore the potential of PrpA protein in promoting IL-10 production by DCs, murine DCs were stimulated with rPrpA protein (10 μg/mL) for 7 hours. Surprisingly, the results indicated that rPrpA protein did not induce IL-10 secretion by DCs (Fig 1E). Previous studies have highlighted PrpA’s ability to promote B lymphocyte proliferation(20, 22). Consequently, we hypothesized that PrpA might also induce murine macrophage proliferation. To validate this hypothesis, macrophages were transfected with pcDNA3.1-*prpA* or vector, and the proliferation of macrophages was assessed using CCK8 kits. The results supported the notion that PrpA protein indeed induces macrophage proliferation (Fig 1F). These findings collectively suggest that *B. melitensis* M5-90 PrpA can induce macrophages to secrete IL-10 and serves as a potent IL-10 inducer in *vivo*. Additionally, PrpA promotes macrophage proliferation.

### PrpA as an Essential Virulence Factor for *B. melitensis* M5-90

Prior studies have demonstrated the importance of IL-10 for the intracellular survival of *Brucella*(18, 21, 28). Given PrpA’s role as a potent IL-10 inducer, it is postulated that PrpA may influence *Brucella*’s intracellular survival by inducing IL-10 in macrophages. To test this assumption, macrophages were exposed to *B. melitensis* M5-90, *B. melitensis* M5-90 *prpA* mutant, or *B. melitensis* M5-90 *prpA*-c (pBBR-prpA). The intracellular survival of the *B. melitensis* M5-90 *prpA* mutant was significantly lower than that of the parental strain, while complementation of *prpA* markedly enhanced *Brucella*’s intracellular survival, even surpassing that of the parental strain (Fig 2A).

**Fig. 2:**
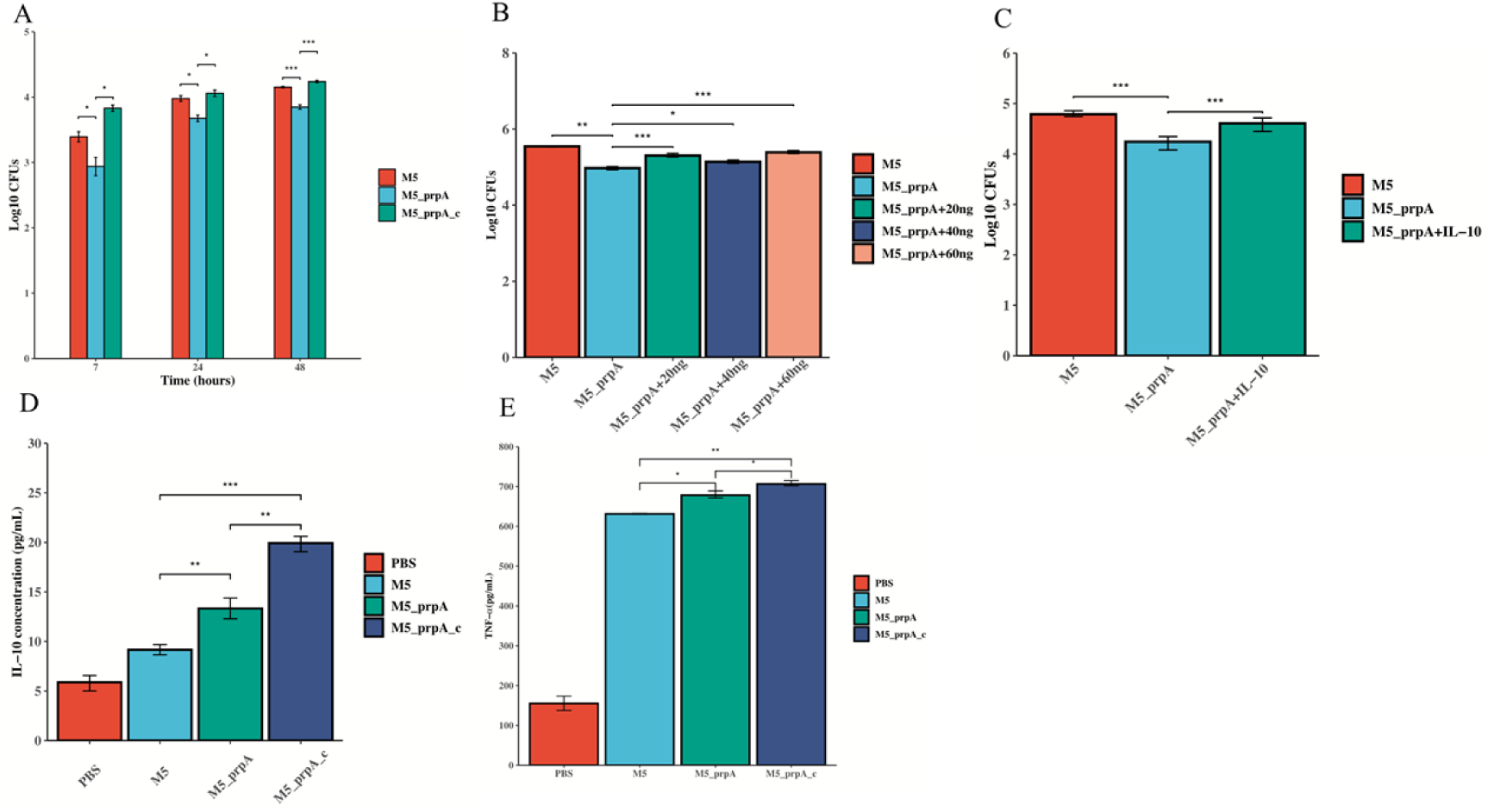
PrpA as an Essential Virulence Factor for *Brucella* Survival. A) Evaluation of intracellular survival of *B. melitensis* M5-90, *B. melitensis* M5-90 *prpA* mutant, and *B. melitensis* M5-90 *prpA* complement strain (*B. melitensis* M5-90 *prpA*_c) in RAW264.7 cells. Macrophages were infected, and CFUs were respectively counted at 7, 24 and 48 hours post-infection (PI). B) Assessment of the impact of varying doses of exogenous IL-10 on intracellular survival of *B. melitensis* M5-90 *prpA* mutant. Macrophages were infected, and 20 ng, 40 ng, and 60 ng of exogenous IL-10 were added to the *B. melitensis* M5-90 *prpA* mutant group. CFUs were counted at 7 h PI. C) Examination of exogenous IL-10’s effect on the survival of *B. melitensis* M5-90 *prpA* mutant in mice. Balb/c female mice were immunized, and 50 μg/kg exogenous IL-10 was injected into the *B. melitensis* M5-90 *prpA* mutant group. CFU numbers in spleen homogenates were counted at 15 days PI. D-E). Measurement of IL-10 and TNF-α levels induced by *B. melitensis* M5-90, *B. melitensis* M5-90 *prpA* mutant, or *B. melitensis* M5-90 *prpA*-c in macrophages.

To further elucidate the impact of IL-10 on *Brucella*’s intracellular survival, macrophages were infected with *B. melitensis* M5-90 or *B. melitensis* M5-90 *prpA* mutant, and varying doses of exogenous IL-10 were added. Results indicated that exogenous IL-10 markedly increased the intracellular survival of the *B. melitensis* M5-90 *prpA* mutant (Fig 2B). Moreover, exogenous IL- 10 also enhanced *Brucella*’s survival in *vivo* (Fig 2C). To investigate whether the deletion of *prpA* affects IL-10 production during infection, cell supernatant was collected when macrophages were infected with *B. melitensis* M5-90, *B. melitensis* M5-90 *prpA* mutant, *B. melitensis* M5-90 *prpA*-c, or PBS (uninfected group). Results demonstrated that the IL-10 level in the *B. melitensis* M5-90 *prpA* mutant group was significantly higher than that of the parental strain. Additionally, complementation of *prpA* further increased IL-10 production in macrophages (Fig 2D). Similar patterns were observed in the levels of TNF-α in the cell supernatant among these groups (Fig 2E). In summary, PrpA serves as a crucial virulence factor, playing a significant role in *Brucella*’s intracellular survival. Furthermore, the deletion of *prpA* significantly enhances *Brucella*’s pro-inflammatory abilities.

### *B. melitensis* M5-90 PrpA Induces Macrophages to Secrete IL-10 via Interaction with Tpl2

Considering the pivotal role of PrpA as a key virulence factor and its impact on *B. melitensis* M5- 90 survival through IL-10 induction in macrophages, it becomes crucial to elucidate the mechanisms by which PrpA promotes IL-10 production in macrophages during infection. Macrophages were infected with *B. melitensis* M5-90 or *B. melitensis* M5-90 *prpA* mutant, and a label-free relative quantitative proteomic method was employed to identify differentially abundant proteins in macrophages. The results revealed a total of 76 differentially expressed (DE) proteins, with 62 being down-regulated and 14 up-regulated (Fig 3A). The majority of these DE proteins were associated with cellular processes, biological regulation, and metabolic processes (S2A Fig). Notably, the down-regulated Tpl2 (Map3k8) protein may be associated with PrpA-induced IL-10 secretion, analogous to its role in *Mycobacteria tuberculosis* infection (29).

**Fig. 3:**
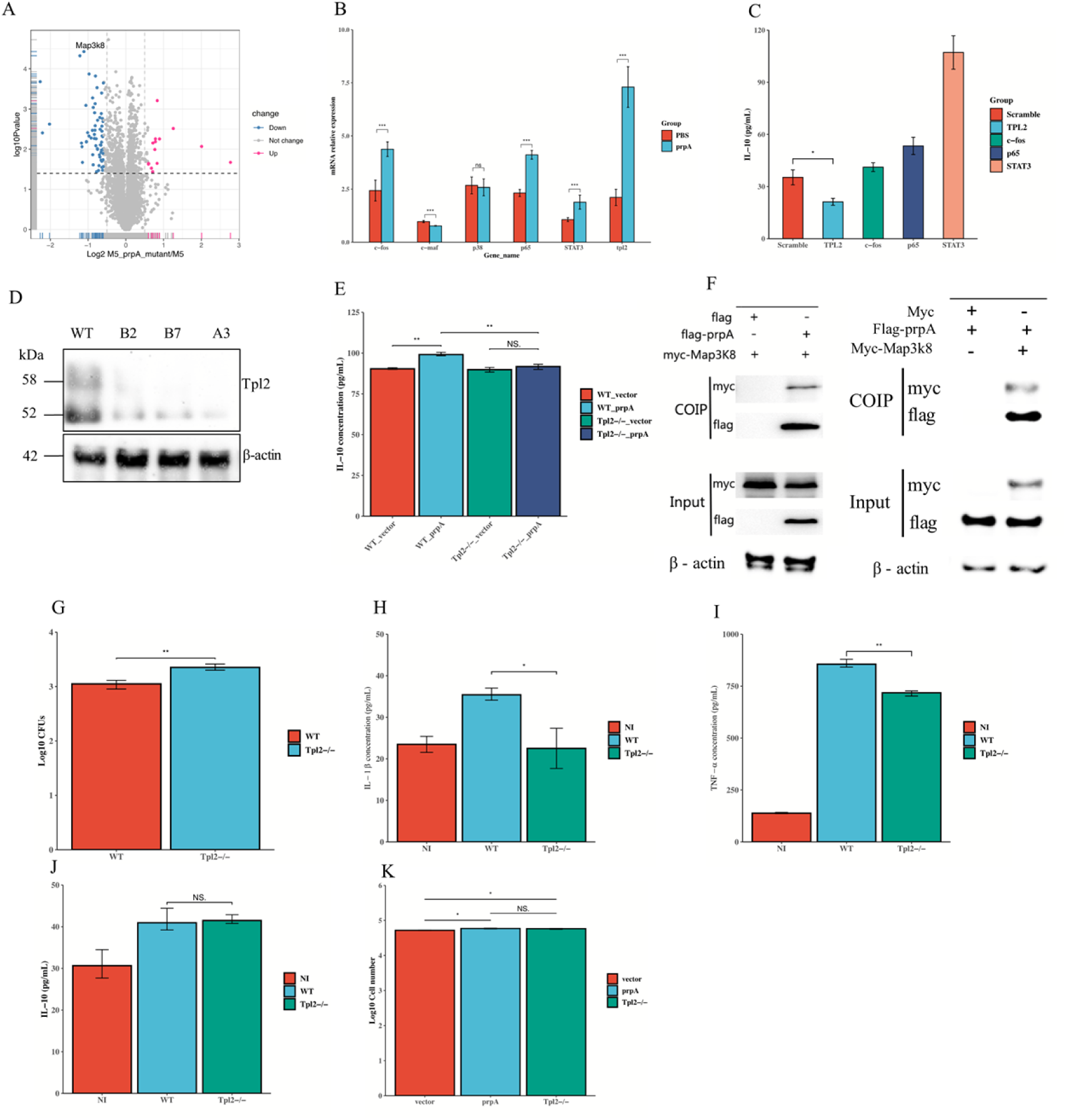
PrpA-Mediated Promotion of IL-10 Production in Macrophages via Tpl2 Interaction during *Brucella* Infection. A) Volcano plot illustrating differentially expressed proteins in macrophages infected with *B. melitensis* M5-90 or *B. melitensis* M5-90 *prpA* mutant. B) Analysis of IL-10 secretion-associated gene expression levels using RT-qPCR after macrophages were stimulated with rPrpA or PBS for 7 h. The expression of *c-fos*, c-maf, p38, *NF-κBp65*, *STAT3*, and *Tpl2* was normalized to *GAPDH* expression. C) Determination of IL-10 concentration induced by rPrpA in macrophages after treatment with siRNA against *Tpl2*, *c-fos*, *NF-κBp65*, and *STAT3*. D) Identification of *Tpl2*^-/-^ macrophages using western blotting. A3 strain, chosen for this study, exhibited complete *Tpl2* knockout via CRISPR-Cas9. E) Measurement of IL-10 concentration induced by PrpA in macrophages after *Tpl2* knockout. F) Analysis of PrpA and Tpl2 interaction by transfecting plasmids into HEK293T cells. G) Enumeration of CFUs in WT or *Tpl2*^-/-^ macrophages infected with *B. melitensis* M5-90 at 24 h PI. H–J. Assessment of the effect of *Tpl2* deletion on IL- 1β, TNF-α, and IL-10 levels in macrophages infected with *B. melitensis* M5-90 at 24 h PI. K) Evaluation of the effect of *Tpl2* deletion on macrophage proliferation induced by pcDNA3.1-*prpA* transfection.

Previous studies have implicated several genes, including *c-fos*(30), *c-maf*(31), *p38*(32), *NF- κB p65*(33), *STAT3*(34), and *Tpl2*(29, 30), in IL-10 secretion in murine macrophages. To investigate whether these genes are involved in PrpA-mediated IL-10 production, macrophages were stimulated with rPrpA for 7 hours, and the expression levels of target genes were assessed using RT-qPCR. The results demonstrated a significant induction in the expression levels of *c-fos*, *NF-κB p65*, *STAT3*, and *Tpl2* in the PrpA group compared to the PBS group (Fig 3B).

To further investigate the contribution of these genes to PrpA-mediated IL-10 production, macrophages were transfected with siRNAs against these genes or scramble siRNA (as a negative control) and then stimulated with rPrpA. The results indicated a marked reduction in the concentration of IL-10 when the expression of *Tpl2* was inhibited with siRNA (Fig 3C). Furthermore, to validate Tpl2’s involvement in PrpA-mediated IL-10 production in macrophages, *Tpl2* was knocked out using the CRISPR-Cas9 approach. The expression of Tpl2 was examined using western blotting (Fig 3D) and RT-qPCR (S2B Fig). The results demonstrated that the deletion of *Tpl2* significantly reduced the IL-10 production induced by stimulation with PrpA (Fig 3E).

### Interaction Between PrpA and Tpl2 in Macrophages for IL-10 Production

Building upon the earlier data, we formulated a hypothesis suggesting an interaction between PrpA and Tpl2 in macrophages leading to IL-10 production. To validate this assumption, HEK293T cells were transfected with pcDNA3.1-*prpA*-Flag, pcDNA3.1-*Tpl2*(*Map3k8*)-Myc, or vector. The results demonstrated a clear interaction between PrpA and Tpl2 (Fig 3F).

Previous research has underscored Tpl2’s critical role in host defense against *Listeria monocytogenes* and *Mycobacterium tuberculosis*. Additionally, *Tpl2* deletion in macrophages has been associated with a significant reduction in IL-1β and TNF-α production(29, 35). However, the role of *Tpl2* in *Brucella* infection remains unknown. To investigate this, wild-type (WT) macrophages or *Tpl2*^-/-^ macrophages were infected with *B. melitensis* M5-90. The results indicated a substantial reduction in the bactericidal ability of macrophages upon *Tpl2* deletion (Fig 3G and S2C Fig). Concurrently, we measured the levels of IL-1β and TNF-α when WT macrophages or *Tpl2*^-/-^ macrophages were infected with *B. melitensis* M5-90, revealing a decrease in the secretion of these two cytokines during *Brucella* infection (Fig 3H and Fig 3I).

Given Tpl2’s involvement in PrpA-mediated IL-10 production (Fig 3C and Fig 3E), we assessed IL-10 levels when WT macrophages or *Tpl2*^-/-^ macrophages were infected with *B. melitensis* M5-90. Surprisingly, the deletion of *Tpl2* did not impact IL-10 production during *Brucella* infection (Fig 3J). As previous data indicated PrpA’s ability to induce macrophage proliferation (Fig 1E), we explored whether *Tpl2* is involved in PrpA-mediated macrophage proliferation. The results revealed that the deletion of *Tpl2* did not affect macrophage proliferation following stimulation with PrpA (Fig 3K).

### Regulation of IL-10 Production and Bactericidal Activity by *Tpl2* in the Context of *B. melitensis* M5-90 PrpA

The KEGG pathway analysis highlights Tpl2’s affiliation with the MAPK pathway. Previous studies propose Tpl2’s regulatory role in TNF-α production through the activation of extracellular signal-regulated kinase (ERK) downstream of TLR4(36, 37). Therefore, we hypothesized that *Tpl2* deletion might diminish ERK expression. To confirm this, pcDNA3.1-*prpA* or vector was transfected into both WT and *Tpl2*^-/-^ macrophages. Data indicated a lower gene expression of ERK in *Tpl2*^-/-^ macrophages compared to WT macrophages (S2D Fig). Further assessment through western blotting confirmed the reduction in ERK and phospho-ERK expression due to *Tpl2* deletion (S2E Fig).

Additionally, KEGG database analysis identified *TLR2*, *TLR4*, *TLR5*, and *RasGRP1* as upstream genes in the MAPK and Toll-like receptor signaling pathways (<u>map04010</u> and <u>map04620</u>) (S2F Fig and S2G Fig). These genes were considered potential contributors to PrpA-induced IL-10 production. Transfection of pcDNA3.1-*prpA* into WT macrophages validated this assumption, showing marked induction of *RasGRP1*, *TLR4*, and *TLR5* expression by PrpA, while *TLR2* expression remained unchanged (S2H Fig – S2K Fig).

In summary, these findings demonstrate that *B. melitensis* M5-90 PrpA induces macrophages to produce IL-10 by interacting with Tpl2 and subsequently activating ERK. Tpl2 plays a crucial role in the bactericidal ability of macrophages and regulates TNF-α and IL-1β production during *Brucella* infection. Notably, *Tpl2* does not participate in the PrpA-mediated proliferation of macrophages. The activation of the Tpl2-ERK signaling pathway by *B. melitensis* M5-90 PrpA may facilitate IL-10 secretion through the induction of *RasGRP1*, *TLR4*, and *TLR5*.

### Identification of Key Amino Acids in *B. melitensis* M5-90 PrpA for Tpl2 Interaction and IL- 10 Production

The interaction between *B. melitensis* M5-90 PrpA and Tpl2, facilitating IL-10 production in macrophages, prompts the exploration of the specific amino acids contributing to this process. Constructing a three-dimensional model of PrpA and Tpl2 using the SWISS-MODEL server unveiled their structural characteristics (Fig 4A, left: PrpA, right: Tpl2). Selection of the optimal templates, based on GMQE scores, revealed similarities of 41.27% and 94.55% for PrpA and Tpl2, respectively. The models exhibited good quality as indicated by GMQE values (PrpA: 0.82, Tpl2: 0.68) and QMEAN scores (PrpA: -0.42, Tpl2: -0.76). Ramachandran plot analysis showed that the majority of the residues of PrpA (94.5%) and Tpl2 (92.3%) in favored regions (S3A Fig and S3B Fig). The Z-score was computed by comparing the normalized raw scores of the PrpA or Tpl2 model (which included the composite QMEAN score and individual mean force potential terms) to a collection of high-resolution X-ray structures present in the PDB database(38). The Z-score of the PrpA protein or Tpl2 protein model indicates that they are located in a relatively accurate region (S3C Fig and S3D Fig). These results confirmed the models’ accuracy for PrpA and Tpl2.

**Fig. 4:**
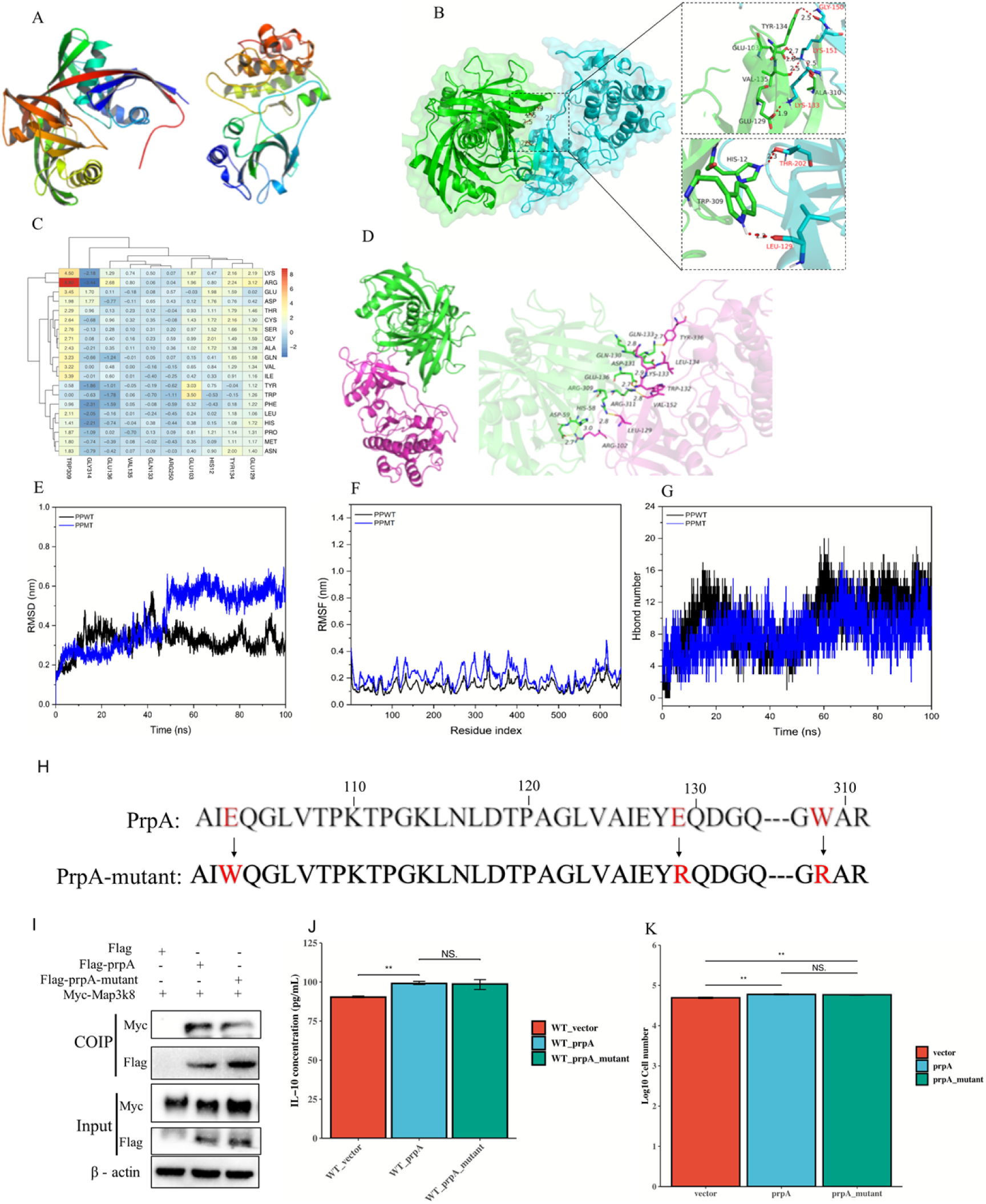
Crucial Residues Trp309, Glu103, and Glu129 in the PrpA-Tpl2 Interaction. A) Protein model depicting PrpA (left) and Tpl2 (right). B) Analysis and interpretation of docking outcomes between PrpA and Tpl2. C) Identification of key interacting residues in the PrpA-Tpl2 interaction using computational mutagenesis. D) Analysis and interpretation of docking outcomes between PrpA-mutant and Tpl2. E) Root Mean Square Deviation (RMSD) analysis of the first 100 ps of molecular dynamics simulation for PrpA-Tpl2 and PrpA-mutant-Tpl2 systems. F) Root Mean Square Fluctuation (RMSF) analysis of molecular dynamics simulation for PrpA-Tpl2 and PrpA-mutant-Tpl2 systems. G) Number of hydrogen bonds formed in the PrpA-Tpl2 and PrpA-mutant- Tpl2 systems during molecular dynamics simulation. H) Model illustrating specific mutated residues in PrpA. I) Analysis of PrpA or PrpA-mutant and Tpl2 interaction after transfection into HEK293T cells. J) Evaluation of the impact of PrpA residue mutations on IL-10 synthesis in RAW264.7 macrophages transfected with pcDNA3.1-*prpA* or pcDNA3.1-*prpA*-mutant plasmids, with pcDNA3.1 vector as a negative control. K) Assessment of the impact of PrpA residue mutations on macrophage proliferation.

ZDOCK simulation predicted the interaction interface between PrpA and Tpl2, emphasizing the involvement of His12, Tyr134, Trp309, and Glu129 through hydrophobic interactions and hydrogen bonds (Fig 4B and S3E Fig). Computational mutagenesis targeting Trp309, Glu103, and Glu129 demonstrated a marked reduction in affinity between PrpA and Tpl2 upon mutation to Arg, Trp, and Arg, respectively (Fig 4C). This substantiates the pivotal role of these amino acids in the PrpA-Tpl2 interaction.

Molecular dynamics simulation confirmed the impact of Trp309, Glu103, and Glu129 mutations on the interaction stability. The mutated PrpA displayed decreased binding energy, increased RMSD values, higher RMSF values, fewer hydrogen bonds, elevated SASA, and expanded Rg compared to the wild-type PrpA (Fig 4D–G and S3F, 3G Fig). These findings collectively indicate that the mutation of these three amino acids compromises the affinity between PrpA and Tpl2.

### Investigation of Amino Acid Mutations in PrpA-Tpl2 Interaction

To explore the impact of mutations in the three critical amino acids on the affinity between PrpA and Tpl2, a PrpA-mutant was generated based on the earlier prediction (Fig 4H) and inserted into the pcDNA3.1-Flag-C vector. HEK293T cells were then transfected with pcDNA3.1-*prpA*, pcDNA3.1-*prpA*-mutant, or the vector alone. Results demonstrated that the mutation of these amino acids slightly weakened the affinity between PrpA and Tpl2 (Fig 4I). Moreover, the effect of these mutations on PrpA-mediated IL-10 production and proliferation in macrophages was evaluated. Surprisingly, the mutation did not influence the PrpA’s ability to induce IL-10 production and proliferation (Fig 4J and Fig 4K). These findings emphasize the dispensable role of Trp309, Glu103, and Glu129 in the PrpA-Tpl2 interaction while highlighting their non-involvement in PrpA- mediated IL-10 production and macrophage proliferation.

### *B. melitensis* M5-90 *prpA* Mutant as a Superior Vaccine Candidate

Considering the potential of *B. melitensis* M5-90 PrpA to induce IL-10 production and suppress the immune response, the deletion of *prpA* in the *B. melitensis* M5-90 may enhance the immune response following immunization. To test this hypothesis, mice were immunized with *B. melitensis* M5-90, *B. melitensis* M5-90 *prpA* mutant, or *B. melitensis* M5-90 *prpA*-c strains. Blood was collected at 30 and 45 days post-immunization (PI), and then the mice was euthanized for spleen CFU enumeration, and mice were challenged with virulent *B. melitensis* M28 at 50 days PI (Fig 5A). Results revealed a lower CFU count in the *B. melitensis* M5-90 *prpA* mutant group compared to the *B. melitensis* M5-90 group at 30 days PI, while the *B. melitensis* M5-90 *prpA*-c group showed a similar CFU count to the *B. melitensis* M5-90 *prpA* mutant group (Fig 5B). By 45 days PI, *Brucella* was cleared in all three groups. Additionally, the mean body weight of mice in the *B. melitensis* M5-90 *prpA* mutant group surpassed that of the *B. melitensis* M5-90 group at both time points, while the *B. melitensis* M5-90 *prpA*-c group exhibited similar weights to the *B. melitensis* M5-90 *prpA* mutant group (Fig 5C). The comparable phenotype between the *B. melitensis* M5-90 *prpA*-c and *B. melitensis* M5-90 *prpA* mutant groups led to the exclusion of the former from subsequent figures.

**Fig. 5:**
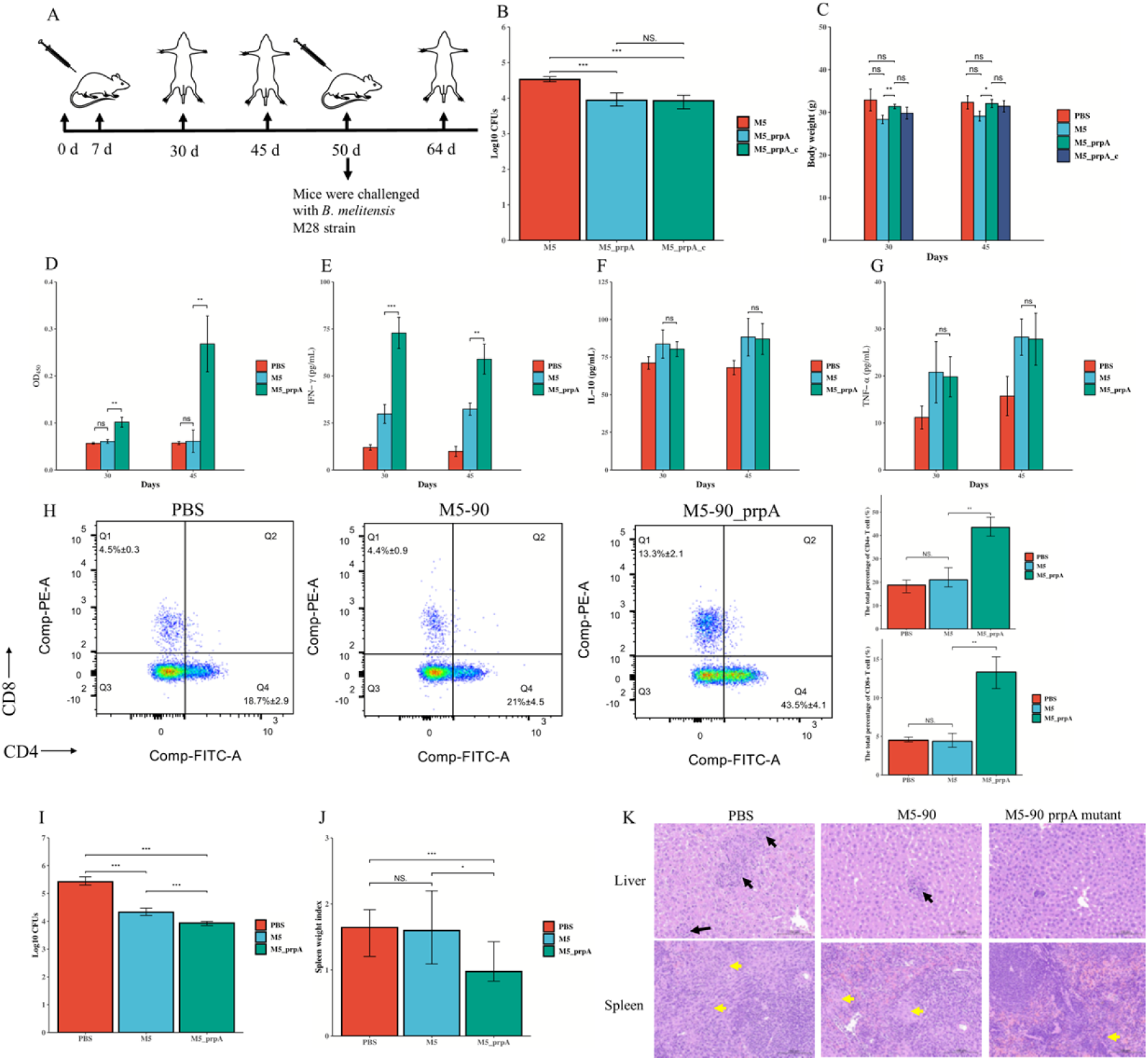
Enhanced Immune Response and Protection Induced by *B. melitensis* M5-90 *prpA* Mutant in Mice. A) Illustration of the immunization and challenge route in mice. B) Enumeration of Colony Forming Units (CFUs) in mouse spleens 30 days post-infection with *B. melitensis* M5- 90, *B. melitensis* M5-90 *prpA* mutant, or *B. melitensis* M5-90 *prpA*_c. C) Measurement of body weight in mice immunized with *B. melitensis* M5-90, *B. melitensis* M5-90 *prpA* mutant, or *B. melitensis* M5-90 *prpA*_c at 30 and 45 days. D-G) Determination of specific anti-*Brucella* IgG, IFN- γ, IL-10, and TNF-α levels in mice post-immunization with *B. melitensis* M5-90, *B. melitensis* M5- 90 *prpA* mutant, or PBS at 30 and 45 days. H) Quantification of CD4+ and CD8+ T cells in mice immunized with *B. melitensis* M5-90, *B. melitensis* M5-90 *prpA* mutant, or PBS at 30 days. I) Enumeration of CFUs in mouse spleens two weeks post-challenge with *B. melitensis* M28 (i.p.). J) Calculation of spleen weight index after mice were challenged with *B. melitensis* M28 for two weeks, presented as organ weight (in grams) per gram of mouse body weight. K) Histopathology of liver and spleen from immunized mice post-challenge with *B. melitensis* M28 for two weeks, with black and yellow arrows indicating microgranulomas.

### Enhanced Immune Response and Protection Induced by *B. melitensis* M5-90 *prpA* Mutant

In order to evaluate the immunogenicity elicited by *B. melitensis* M5-90 *prpA* mutant in comparison to *B. melitensis* M5-90, the levels of IFN-γ and specific anti-*Brucella* IgG in the serum were assessed at 30 and 45 days post-immunization (PI). Results indicated elevated levels of specific anti-*Brucella* IgG and IFN-γ in the *B. melitensis* M5-90 *prpA* mutant group compared to the *B. melitensis* M5-90 group at both time points (Fig 5D and Fig 5E). Notably, similar levels of IL-10 and TNF-α were observed in both groups (Fig 5F and Fig 5G).

Previous studies have underscored the significant role of CD4+ T cells in IFN-γ production during adaptive immune responses, especially in the context of *Mycobacterium tuberculosis* infection(39, 40). Furthermore, IFN-γ is crucial for orchestrating an effective CD8+ T cell response. Given the markedly higher IFN-γ levels induced by the *B. melitensis* M5-90 *prpA* mutant (Fig 5E), we hypothesize an enhanced influx of both CD4+ and CD8+ T cells. To validate this hypothesis, splenic lymphocytes were isolated from mice vaccinated with either *B. melitensis* M5-90 or *B. melitensis* M5-90 *prpA* mutant. Results revealed an increased count of CD4+ and CD8+ T cells in the *B. melitensis* M5-90 *prpA* mutant group after 30 days PI, although no significant disparity was observed after 45 days PI (Fig 5H and S4 Fig). To further investigate whether IFN-γ is released by the CD4+ and CD8+ T cells, isolated splenic lymphocytes were stimulated with Con A (1 μg/mL) and Brefeldin A (5 μg/mL) and subsequently labeled with FITC anti-mouse CD4+ and PerCP anti-mouse IFN-γ or PE anti-mouse CD8a and PerCP anti-mouse IFN-γ antibodies. The results indicated that no signal was detected in the CD4+ or CD8+ T cells gates for both in *B. melitensis* M5-90 and *B. melitensis* M5-90 *prpA* mutant groups at 30 or 45 days PI. Therefore, the elevated level of the IFN-γ in the *B. melitensis* M5-90 *prpA* mutant group may be attributed to secretion by other immune cells.

Although PrpA is recognized to promote B lymphocyte proliferation(20), it remain unclear whether the elevated percentage of CD4+ and CD8+ T cells is attributable to the reduction in B cells or an increase in T cell numbers. To address this hypothesis, T cell counts were quantified in isolated splenic lymphocytes. The results demonstrated no significant difference in T cell numbers between the *B. melitensis* M5-90 and *B. melitensis* M5-90 *prpA* mutant groups at 30 or 45 days PI (S5 Fig). Thus, the deletion of *prpA* in *B. melitensis* M5-90 led to a reduction in B lymphocytes during vaccination.

The data suggests that the absence of *prpA* enhances the immune response against *Brucella*, prompting the hypothesis that *B. melitensis* M5-90 *prpA* mutant may confer superior protection compared to *B. melitensis* M5-90. In-depth analysis involved immunizing mice with either strain and challenging them with virulent *B. melitensis* M28 at 50 days PI. Subsequent isolation of spleen and liver two weeks post-challenge revealed significantly diminished CFUs in the *B. melitensis* M5- 90 *prpA* mutant group compared to the *B. melitensis* M5-90 group (Fig 5I). Additionally, the spleen weight index was lower in the *B. melitensis* M5-90 *prpA* mutant group (Fig 5J), and a notable reduction in granuloma formation within the liver and spleen was observed (Fig 5K). Overall, these findings suggest that PrpA is a pivotal virulence factor for *Brucella* survival in mice, and the deletion of *prpA* augments the immune response, providing higher protection against wild-type strain infection.

## Discussion

IL-10, acknowledged as an anti-inflammatory cytokine, assumes a pivotal role in thwarting inflammatory and autoimmune disorders. It is noteworthy that various pathogens employ diverse strategies to hinder or circumvent immune responses by inducing immune cells, including macrophages, T cells, B cells, neutrophils, and certain dendritic cells, to generate IL-10(41).

Pathogens such as *Leishmania major*(42), *Human cytomegalovirus*(43), *Mycobacteria tuberculosis*(44), *Salmonella*(17), and *Listeria monocytogenes*(45) exploit IL-10 to prolong their persistence and establish chronic infections. Moreover, numerous studies have substantiated that *Brucella* can adeptly modulate immune responses by instigating immune cells to secrete IL-10, thereby establishing persistent infection(18, 21, 28).

To date, only *B. abortus* 2308 PrpA has been identified as a potent IL-10 inducer, interacting with the surface receptor NMM-IIA to facilitate IL-10 production while restraining macrophages from secreting proinflammatory cytokines(20, 23). To explore the possibility of other virulence factors in *Brucella* inducing IL-10 production, three additional factors (OMP25, wadC, and RomA) were selected due to their deletion resulting in heightened immune responses during infection. However, solely PrpA emerged as an IL-10 inducer both in *vitro* and in *vivo*. Consequently, the modulation of the immune response by the other three virulence factors may occur through alternative pathways during infection.

Interestingly, PrpA exhibited the capacity to stimulate IL-10 production in macrophages and mice but not in DCs, although most of the innate receptors of DCs and macrophages are common to both these types of cell. Extant research has elucidated a stark dichotomy in the responses of DCs and macrophages to LPS. DCs upon LPS exposure, acquire migratory capacities, secretory inflammatory cytokines, undergo terminal differentiation, and ultimately succumb to apoptosis (46). Conversely, LPS-activated macrophages initiate inflammation but subsequently persist in tissue, transitioning to an anti-inflammatory phenotype to facilitate homeostasis restoration (47, 48). Hence, it is plausible that DCs and macrophages may likewise exhibit distinct response patterns to PrpA. In addition, the macrophages and naïve splenocytes represent the primary targets of PrpA, inducing these cell types to produce IL-10(20, 23), it can be inferred that these cells constitute the primary source of IL-10 when stimulated with rPrpA in mice. However, the potential of PrpA to target other cell types remains an unresolved aspect.

Tpl2, a serine-threonine kinase, assumes a critical role in the innate immune response by bridging toll-like receptors (TLRs) and TNF production through ERK activation(36). Previous investigations underscored Tpl2’s significance in host defense against *Listeria monocytogenes* infection, where *Tpl2*-deficient mice exhibited elevated pathogen burden compared to wild-type counterparts. *Tpl2* deletion not only reduced IL-1β production in macrophages and DCs but also left TNF production unimpaired(35). Furthermore, Tpl2’s involvement in host defense against *Mycobacterium tuberculosis* has been elucidated, with *Tpl2*-deficient mice displaying heightened susceptibility to the pathogen. *Tpl2* deletion resulted in the abrogation of ERK activation, subsequently reducing IL-10, TNF-α, and IL-1β production(29, 49). In the current study, *Tpl2* deletion significantly compromised macrophage bactericidal ability during *Brucella* infection, accompanied by decreased IL-1β and TNF-α production. This reduction may contribute to the higher colony-forming units (CFUs) in *Tpl2*-deficient macrophages compared to wild-type counterparts, as IL-1β and TNF-α are crucial pro-inflammatory cytokines in the host’s defense against *Brucella* infection. Surprisingly, the IL-10 level remained unimpaired in *Tpl2*-deficient macrophages during *Brucella* infection, suggesting that *Brucella*, may target multiple pathways in macrophages to induce IL-10, and other virulence factors in *Brucella* may also induce macrophages to produce IL-10. Additionally, besides *Tpl2*, four other genes (*c-fos*, *c-maf*, *NF-κB p65*, and *STAT3*) were identified as PrpA targets, although their associated mechanisms were not explored here. Notably, *c-fos* has been implicated in proper cell division during *Brucella* infection and efficient restriction of *Brucella* invasion and survival in macrophages(50). This hints at a potential role of *c-fos* in PrpA-mediated macrophage proliferation. Furthermore, our study identified *RasGRP1*, an upstream gene in the Tpl2-ERK signaling pathway, as induced by PrpA. *RasGRP1* has been associated with the promotion of acute inflammation and inhibition of inflammation-associated cancer(51). While PrpA did not induce TNF-α production in macrophages, it induced *RasGRP1* expression, suggesting that PrpA may trigger the secretion of other pro-inflammatory cytokines through *RasGRP1*.

Moreover, TLR2, TLR4, and TLR5 play crucial roles in pathogen recognition(52, 53). PrpA induced the expression of *TLR4* and *TLR5*, implying a potential interaction between PrpA and these TLRs. This raises the hypothesis that *TLR4* and *TLR5* might serve as new receptors for PrpA in inducing IL-10 production in macrophages, warranting further investigation.

A previous study has indicated that the polyclonal activation of lymphocytes induced by pathogens can be detrimental to the host’s immune response(54). Experimental evidence demonstrates that PrpA facilitates the proliferation of B lymphocytes, contingent upon the intactness of the proline-racemase active site(20). Moreover, our current study reveals that PrpA induces the proliferation of macrophages. However, the role of macrophage proliferation during infection is complex. Prior research has shown a correlation between the expansion of resident alveolar macrophages and resistance to *L. sigmodontis* infection in C57Bl/6 mice, contrasting with Balb/c mice, which exhibit susceptibility associated with the accumulation of infiltrating macrophages(55). The proliferation of macrophages has been linked to cellular resistance against *Listeria monocytogenes* infection(56). Recent findings suggest that *ccnd1*/*ccnd2* regulate macrophage proliferation during *Brucella* infection, but this proliferation does not impact the host’s defense against *Brucella*(50). The mitogenic activity of PrpA, previously shown to depend on Cys253, although this alone is not adequate for its activity(20). Hence, we examined the other three amino acids (Trp309, Glu103, and Glu129) of PrpA in this study, however, with no influence observed from the mutation of these three other residues on macrophage proliferation or IL-10 production. Notably, Cys8 and Cys14 were found to be essential for S-palmitoylation, crucial for PrpA’s immunomodulatory functions(23). However, the involvement of Cys8, Cys14, and Cys253 in PrpA- mediated IL-10 production or interaction with Tpl2 remains unknown, representing a limitation of this study.

In a previous investigation, the deletion of *prpA* led to decreased survival of *Brucella* in mice, although invasion and intracellular replication in macrophages showed no significant variances between the *prpA* mutant strain of *B. abortus* 2308 and its parental strain(20). Conversely, in our study, the *B. melitensis* M5-90 *prpA* mutant exhibited an attenuated phenotype both in *vitro* and in *vivo*. This discrepancy could arise from distinct pathogenic mechanisms between the wild strain and the vaccine strain of *Brucella*(57). Furthermore, complementing *prpA* in *B. melitensis* M5-90 *prpA* mutant did not restore its survival in mice, possibly due to the instability of pBBR1-MCS plasmids in mice(58).

Activation of antigen-presenting cells crucial for *Brucella* eradication depends on the cytokines IFN-γ and TNF-α(59, 60). In our study, the serum IFN-γ level from the *B. melitensis* M5-90 group was lower than that in the *B. melitensis* M5-90 *prpA* mutant group at both 30 and 45 days post-infection (PI), consistent with a previous study(22). TNF-α levels were similar between the two groups at both time points, unlike the higher TNF-α level induced by the *B. abortus* 2308 *prpA* mutant compared to its parental strain in mice(22). This discrepancy may stem from different pathogenesis between *B. abortus* and *B. melitensis* in mice(61), warranting further investigation into the underlying mechanisms.

CD4+ and CD8+ T cells play pivotal roles in orchestrating the immune response against intracellular pathogens(62). The IFN-γ produced by these T cells and NK cells activates macrophages’ bactericidal properties, crucial for impeding *Brucella* infection(63–66). Our study uncovers a notable finding that the *B. melitensis* M5-90 *prpA* mutant elicits an elevated abundance of CD4+ and CD8+ T cells compared to its parental strain. This phenomenon is not attributable to an increase in total T cells, but rather associated with PrpA’s role in inducing B lymphocyte proliferation. Notably, B cells can provide an infectious niche for *Brucella*, contributing to the establishment of chronical infection(67, 68). This observation lends further support the notion that PrpA acts as an immune suppression factor during *Brucella* infection, and its deletion significantly enhances the immune responses in mice.

The protection conferred by the *B. melitensis* M5-90 *prpA* mutant surpassed that of the parental strain, as demonstrated in challenge experiments in mice. While these findings underscore the potential of PrpA as a subunit vaccine for brucellosis prevention, it’s noteworthy that PrpA, despite stimulating peripheral blood mononuclear cells to produce IFN-γ, induces lower levels of IgG antibodies compared to other immunogens(66, 69–71). The deletion of *prpA* in our study resulted in a significant elevation of specific anti-*Brucella* IgG levels. This phenomenon may be attributed to the unique character of *Trypanosoma cruzi* proline racemase, homologous to *Brucella* proline racemase, which hinders humoral immune responses through B cell polyclonal activation(72). These results highlight PrpA as a key virulence factor in *Brucella*, modulating the immune response during infection. Future work should focus on understanding the role of specific amino acids in PrpA’s immune suppression function through deletion or mutation, aiming to improve vaccine immunogenicity and protection.

While the *B. melitensis* M5-90 *prpA* mutant demonstrated a higher immune response and protection than the parental strain in mice, the duration of the vaccine’s efficacy raises concerns. Our study indicates a more rapid clearance of the *B. melitensis* M5-90 *prpA* mutant compared to *B. melitensis* M5-90. The correlation between vaccine strain persistence and protection against infection, as established in previous research(73), underscores the need for further evaluation of *B. melitensis* M5-90 *prpA* mutant in cattle or sheep. Assessments should encompass immunogenicity, safety, duration, and overall protective efficacy to ascertain its potential as a viable vaccine candidate.

In this study, five genes (*c-fos*, *c-maf*, *NF-κBp65*, *STAT3*, and *Tpl2*) were discerned as PrpA targets. PrpA’s interaction with Tpl2 activates ERK, facilitating IL-10 production and thereby promoting persistent infection (Fig 6). Notably, three residues (Trp309, Glu103, and Glu129) in PrpA are implicated in the interaction with Tpl2. Furthermore, PrpA may exploit *TLR4*, *TLR5*, and *RasGRP1* to activate the Tpl2-ERK signaling pathway. Deletion of *prpA* adversely impacted the survival of *B. melitensis* M5-90, concurrently enhancing specific anti-*Brucella* IgG, IFN-γ, and the number of CD4+ and CD8+ T cells. Our results collectively unveil the mechanisms by which *Brucella* PrpA induces macrophages to produce IL-10, facilitating the establishment of chronic infection. Moreover, the *B. melitensis* M5-90 *prpA* mutant emerges as a promising vaccine, providing superior protection compared to the parental strain in mice.

**Fig. 6:**
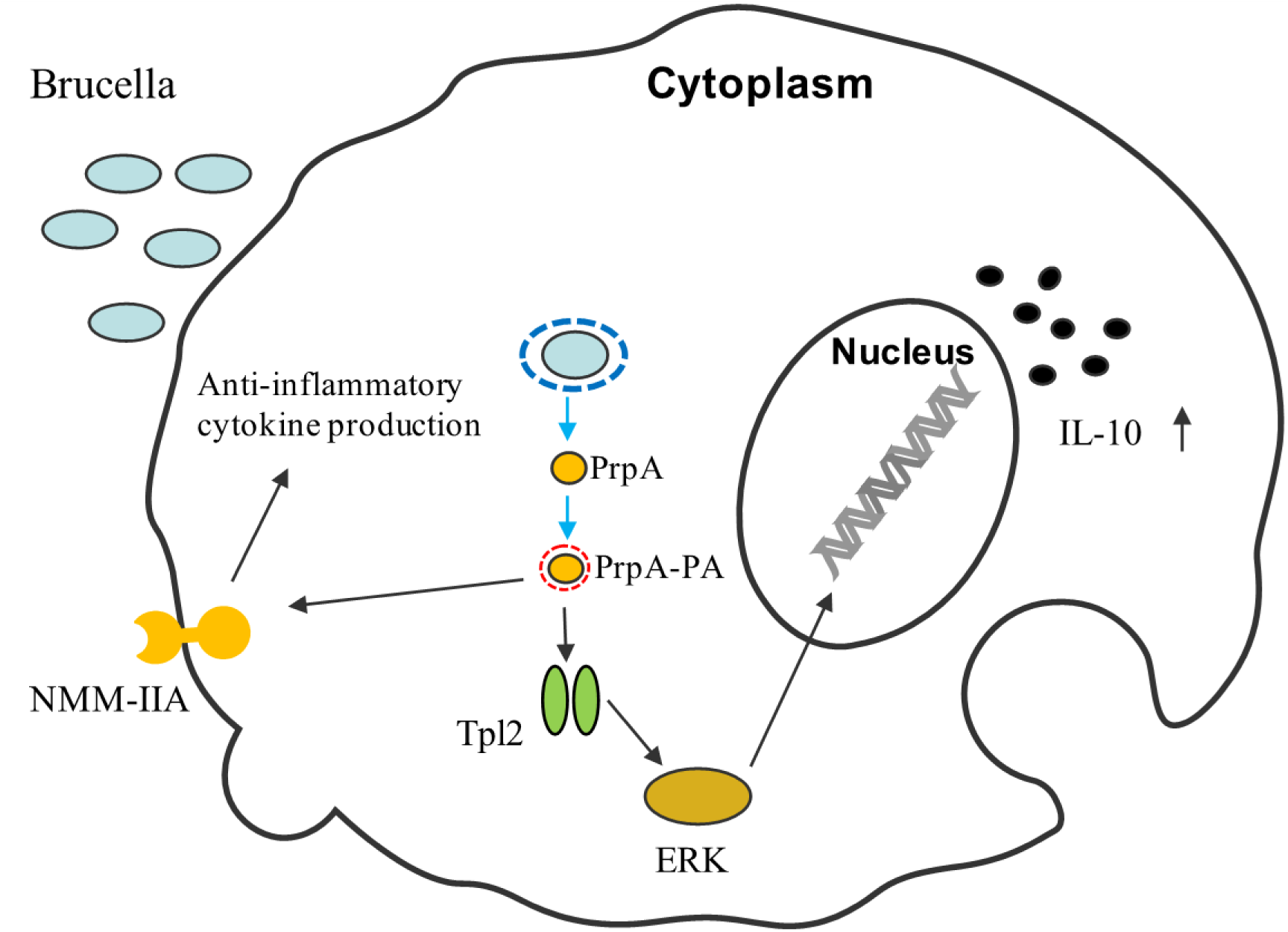
Proposed Model of PrpA and Tpl2 Interaction in Macrophages. The model postulates the interaction between PrpA and Tpl2 in macrophages. PrpA is secreted by *Brucella* into the cytoplasm of infected macrophages, undergoes palmitoylation by the host cell, and is subsequently transported to the plasma membrane. At the plasma membrane, PrpA interacts with the surface receptor NMM-IIA, as previously demonstrated in a prior study(23). In this investigation, we hypothesize that PrpA undergoes palmitoylation initially and subsequently interacts with Tpl2. This interaction, in turn, activates the ERK pathway. The activation of the Tpl2- ERK pathway culminates in the production of IL-10 in macrophages.

## Methods

### Ethics Statement

The care, handling, and experimental procedures involving mice strictly adhered to the regulations and guidelines established by the Animal Care and Use Committees of Shihezi University, as approved for this study.

### Bacterial Strains and Culture Conditions

BL21(DE3) (Invitrogen, USA) was employed for the expression of the outer membrane protein 25 (OMP25), proline-racemase protein A (PrpA), glycosyltransferase (wadC), and periplasmic protein (RomA). Bacterial strains were cultivated at 37°C in Luria-Bertani (LB) broth or agar (Difco Dickinson). *PrpA* deletion mutants of *B. melitensis* M5-90 were created using homologous recombination methods as previously described(74). In brief, *Brucella* was cultured in tryptic soy broth (TSB) (Difco, BD, USA) at 37°C and 150 rpm for 40 h at the exponential stage, made electrocompetent using conventional microbiological techniques. Subsequently, the electrocompetent *Brucella* strains underwent electroporation with the pGEM-7Zf+ plasmid, and *prpA* deletion mutants were identified and isolated in the presence of 100 μg/mL kanamycin. The primers for constructing the *B. melitensis* M5-90 *prpA* mutant are listed in S1 Table. The complementation plasmid for *prpA* was created by PCR amplification of *prpA* ORF, cloned into the pBBR1-MCS4 vector, known for its effectiveness in complementation analysis in *B. melitensis*(75). After digestion, the PCR product was ligated with pBBR1-MCS4, which had also undergone digestion, facilitating the generation of the complementation plasmid. To ensure transcription originating from the lac promoter within the plasmid, the *prpA* amplified product was cloned directionally into pBBR1-MCS4. Subsequently, the resulting complementation plasmid, named pBBR-prpA, was introduced into the *prpA* deletion mutant and selected based on its resistance to ampicillin.

### Production, Purification, and Identification of Recombinant Proteins

Cloning, expression, and purification of recombinant proteins were detailed previously(76). In summary, target genes were PCR-amplified from genomic DNA of *B. melitensis* M5-90 (S2 Table). The PCR products underwent sequencing for nucleotide sequence confirmation. The DNA fragments were cloned into a pET32a+ vector (Novagen, Madison, WI, USA). Thirty milliliters of the culture were inoculated into 1 L of LB broth supplemented with 250 μg ampicillin. After culturing at 220 rpm, 37°C for 3 h, 1 mM isopropyl-β-D-thiogalactopyranoside (IPTG; Amresco, USA) induced recombinant protein expression, and the culture was further incubated for 7 h under the previously described conditions. Cells were harvested by centrifugation at 4,400 g for 15 min. The pellets were suspended in 25 mL of binding buffer (20 mM Tris–HCl, 8 M urea, 500 mM NaCl, 20 mM imidazole, 1 mM β-mercaptoethanol, pH 8.0), sonicated at 10,000 Hz in ice water (50% pulse, 25 s pulse/50 s steps, 20 cycles), and centrifuged at 4,400 g for 10 min to collect supernatants. Recombinant proteins were purified using a His SpinTrap (GE Healthcare, UK) following the manufacturer’s protocol. The concentration of purified recombinant proteins was measured using a BCA kit (Bio-Rad, USA). The purity and identity of the recombinant proteins were analyzed by SDS-PAGE and western blotting with an anti-His antibody (Abcam, UK). Endotoxin contamination in the purified proteins was eliminated by 0.1% Triton X-114 in washing buffer in the purification process and further confirmed using an endotoxin assay kit (Toxin Sensor™ Chromogenic LAL endotoxin Assay kit, GenScript)(77).

### Cells, Culture Conditions, Stimulation, and Cytokine Detection

The murine macrophage RAW264.7 cell line (ATCC, TIB-71), HEK293T (ATCC, CRL-3216), and DC2.4 (Procell, Wuhan) were cultured in Dulbecco’s modified Eagle’s medium (DMEM, Gibco Life Technologies, Rockville, MD, USA) supplemented with 10% fetal bovine serum (Gibco, USA), 100 U/mL penicillin, and 100 μg/mL streptomycin at 37 °C with 5% CO_2_. RAW264.7 and DC2.4 were adjusted to 1 × 10^6^ cells/well in a 6-well plate and stimulated with 10 μg/mL of the four different recombinant proteins for 7 h. The concentration of IL-10 in the cell culture supernatant was determined using ELISA Quantikine Mouse kit (R&D Systems, Minneapolis, MN, USA). Concanavalin A (ConA, 1 μg/mL) was used as a positive control.

### Mice Immunization with Recombinant Proteins and Cytokine Detection

Four to six-week-old female Balb/c mice were obtained from the Experimental Animal Center of the Academy of Military Medical Science (Beijing, China) and had ad libitum access to food and water. An acclimatization period of one week preceded experimentation. Mice were randomly divided into 5 groups (n = 5 per group). Recombinant proteins, namely rOMP25, rPrpA, rwadC, and rRomA, were emulsified in an oil-in-water mixture using Freund’s complete adjuvant (MERK, USA) during the initial injection or Freund’s incomplete adjuvant (MERK, USA) during subsequent injections. The control group received an equal volume of mixture of phosphate buffer saline (PBS, Solarbio) and Freund’s complete or incomplete adjuvant. Mice were immunized subcutaneously with a dosage of 50 μg of purified proteins during each injection, with a two-week time interval between administrations. Serum for cytokine detection was obtained by retro-orbital bleeding under anesthesia at 15 and 30 days after the second immunization, and IL-10 levels were measured using ELISA Quantikine Mouse kit (R&D Systems, Minneapolis, MN, USA).

### Eukaryotic Expression Vectors

*PrpA* was amplified from the plasmid pBBR1MCS4-Flag-prpA, using primers incorporating the EcoRI and XhoI restriction sites in the forward and reverse primers, respectively. PCR products were treated with these restriction enzymes for subsequent insertion into the corresponding restriction sites of pcDNA3.1-Flag-C. Additionally, the tumor progression locus 2 (Tpl2) was amplified from the genome of RAW264.7 cells, and the primers used for amplifying *Tpl2* had the EcoRI and XhoI restriction sites in the forward and reverse primers, respectively. PCR products were cleaved using the specified restriction enzymes for subsequent insertion into the identical restriction sites of pCMV-Myc. Detailed primer sequences are provided in S3 Table.

### Plasmid Transfection

For plasmid transfections, RAW264.7 cells were seeded on 6-well tissue culture plates at 0.6 × 10^6^ cells/well. 3.5 μg of pcDNA3.1-vector or pcDNA3.1-*prpA* plasmid (endotoxin-free) was transfected into cells using Lipo8000 reagent (Beyotime, Shanghai, China) following the manufacturer’s protocol. Twenty-four hours post-transfection, the medium was replaced with fresh medium. Subsequently, at forty-eight hours post-transfection, the cell supernatant was collected, and the cells were subjected to RNA or protein isolation procedures. The concentration of IL-10 in the cell supernatant was determined using ELISA Quantikine Mouse kit (R&D Systems, Minneapolis, MN, USA).

### Cell Proliferation Assay

The effect of PrpA on RAW264.7 cell proliferation was assessed using Cell Counting Kit-8 (CCK- 8). RAW264.7 cells were seeded into 96-well plates and transfected with 200 ng plasmid of pcDNA3.1-vector or pcDNA3.1-*prpA*. One day after transfection, the medium was replaced with fresh medium, and CCK8 solution was added and cultured under 5% CO_2_ at 37°C for 3 h. Subsequently, the absorbance value was measured using a microplate absorbance reader at 450 nm. The standard curve was constructed in accordance with the protocols provided by the CCK-8 kit manufacture, and subsequent cell quantification was performed by using this standard curve.

### Infections and CFUs Assays

RAW264.7 cells were seeded onto 6-well tissue culture plates at 1 × 10^6^ cells per well in a 2 mL culture medium. Infections were carried out with a multiplicity of infection of 100:1, where bacteria were applied to macrophages through centrifugation at 170 g for 10 minutes at 4°C. Subsequently, the cells were incubated for 60 minutes at 37°C under a 5% CO_2_ atmosphere. To eliminate extracellular bacteria, the cells underwent thorough washing with PBS and were then incubated for an additional 60 minutes in medium containing 50 μg/mL gentamicin. Following this, the medium was replaced with fresh medium supplemented with 10 μg/mL gentamicin after one hour. At 7 h, 24 h and 48 h post-infection (PI), the cells were scraped and lysed in PBS/0.1% X-100 Triton (Sigma-Aldrich). The colony-forming units (CFUs) were quantified by plating different dilutions onto tryptic soy agar (TSA) plates.

### Proteomics Analysis

RAW264.7 cells were infected with *B. melitensis* M5-90 or *B. melitensis* M5-90 *prpA* mutant for 24 h. The cells were scraped and sonicated three times on ice using a high-intensity ultrasonic processor (Scientz) in lysis buffer (8M urea, 1% protease inhibitor cocktail). Debris was removed by centrifugation at 12,000 × g at 4°C for 15 min. Subsequently, the supernatant was collected, and the protein concentration was determined with a BCA kit following the manufacturer’s protocol. Then, the protein solution was digested, dissolved, and subjected to capillary electrophoresis followed by timsTOF Pro (Bruker Daltonics) mass spectrometry. The detailed process has been described previously(57). The obtained MS/MS data were processed using Maxquant (v1.6.15.0) and searched against the SwissProt Mus musculus protein database (version 2020.12, 17,063 sequences) concatenated with the reverse decoy database. Protein’s Gene Ontology (GO) function annotation was completed using the InterProScan software (v.5.14-53.0), according to the protein sequence alignment method. Furthermore, annotation of the protein pathway was completed using the Kyoto Encyclopedia of Genes and Genomes (KEGG) database. Significance was attributed to the functions and pathways that exhibited a P-value of less than 0.05, as determined by the two-tailed Fisher’s exact test.

### RNA Extraction and RT-qPCR Analysis

Monolayers of RAW264.7 cells (approximately 0.6 × 10^6^ cells/well) were stimulated with rPrpA (10 μg/mL) for 7 h, and total RNA was extracted from these cells using TRIzol reagent, following the instructions (CWBIO, Beijing, China). DNA removal from the samples was accomplished using TURBO DNA-free (Ambion). Subsequently, the concentration and quality of the samples were assessed using a Nanodrop 2000 spectrophotometer (Thermo, USA). A total of 2 μg of RNA was employed to synthesize cDNA using the HiFiScript cDNA Kit (CWBIO, Beijing, China). Real-time PCR analysis was performed on a ThermoFisher QuantStudio 3 RT PCR-well Q3 instrument (Thermo Fisher, USA) using SYBR Green dye (CWBIO, Beijing, China). The primers for target genes are listed in S4 Table. The real-time PCR conditions were as follows: an initial denaturation step at 95°C for 5 min, followed by 45 cycles of denaturation at 95°C for 30 s and annealing at either 57°C or 60°C for 30 s. The fold changes of each gene were determined using the 2^-ΔΔCT^ method, and the mRNA levels were normalized by comparing them to the expression of *GAPDH*.

### siRNA Transfection

Small interfering RNAs (siRNAs) against mouse *c-fos*, *NF-κBp65*, *STAT3*, *Tpl2* were designed by GENERAL BIOL Co., Ltd. (Anhui, China). Specific siRNAs or negative control (NC) siRNA were transfected into RAW264.7 cells using Lipo8000 (Beyotime, Shanghai, China) according to the manufacturer’s instructions. Gene knockdown was assessed by reverse transcription quantitative PCR (RT-qPCR). All siRNAs used are described in S5 Table.

### CRISPR-Cas9-Mediated Murine *Tpl2* (*Map3k8*) Knockout in RAW264.7 Cells

The *Tpl2* gene knockout plasmid (pLenti-MAP3K8-sgRNA, catalog number L15930) was procured from Beyotime company (Shanghai, China). Subsequently, the plasmid was introduced into competent DH5α cells through a transformation process. In brief, a 1.5 mL sterilized tube containing 20 μL of competent DH5α cells was used to combine a total of 2 μg of plasmid, which was then gently mixed. The mixture was subjected to a temperature of 42°C for 90 s, followed by 5 min on ice. Subsequently, 1 mL of LB medium was added to the tube, and the culture was incubated at 220 rpm, 37°C for 1 h. The competent DH5α cells were collected by centrifugation at 4,400 g for 5 min, and the supernatant was discarded. Finally, the cells were resuspended in 100 μL of sterilized PBS and inoculated onto LB agar plates supplemented with ampicillin (0.25 μg/μL). The suspension was evenly spread using a loop and incubated at 37°C for 24 h. The single colonies were confirmed by sequencing analysis (Sangon Biotech). The recombinant plasmids were extracted with PureLinkTM HiPure kits (Invitrogen). A total of 3.5 μg of the recombinant plasmids was transfected into RAW264.7 cells using Lipo8000, and pLenti-Control-sgRNA (Beyotime, catalog number L00011) was used as the negative control. 4 μg/mL puromycin (Beyotime, catalog number ST551) was added to the cell cultures to select the knockout cells 48 h later. After approximately 10 days, positive cells were acquired through a selection process and subsequently subcloned into 96-well plates to facilitate individual clone growth. These clones were then preserved as cell stocks. Ultimately, verification of the positive cells was conducted using RT-qPCR and western blotting.

### Coimmunoprecipitation and Western Blotting

HEK293T cells cultured in 60-mm-diameter dishes (Thermo Fisher) were transfected with the specified plasmids using Lipo8000. After 48 h, the cells were lysed with lysis buffer (composed of 150 mM NaCl, 50 mM Tris-HCl [pH 7.4], 1% Nonidet P-40, 0.5% Triton X-100, 1 mM EDTA, 0.1% sodium deoxycholate, 1 mM dithiothreitol, 0.2 mM phenylmethylsulfonyl fluoride, and a protease inhibitor cocktail [Sigma-Aldrich]) on ice for 15 min. The supernatant of the cell lysate was collected by centrifugation and subsequently subjected to a preclearing step by incubating with protein G/protein A agarose for 1 hour at 4°C. Following this, the supernatant was incubated with the specified antibodies overnight at 4°C and then precipitated using protein G/protein A agarose for 30 min at room temperature. Subsequently, the precipitated protein G/protein A agarose was centrifuged at 2,000 × g for 10 s and washed three times with PBS. Finally, the proteins bound to the agarose beads were eluted by boiling for 10 min in 2 × loading buffer. The eluted proteins were then subjected to SDS-PAGE and western blotting. The immunoreactive bands were visualized using enhanced chemiluminescence (ECL) reagents (Thermo Fisher).

### 3D Structure Prediction, Validation, and Molecular Docking

The amino acid sequences of PrpA (NCBI accession: WP_002964871) and Tpl2 (NCBI accession: NP_031772) were submitted to the <u>SWISS-MODEL</u> server to search for the best template to build a 3D model. The generated model was downloaded in PDB format and visualized using Pymol v2.1. The quality of the model was examined using <u>ProSA</u> and the <u>PROCHECK</u> server. The structures of both proteins were subjected to the Clean Protein protocol, involving the addition of missing loops and refinement through minimization utilizing the CHARMM forcefield in BIOVIA Discovery Studio (DS) Visualizer v3.5. The resulting minimized structures were then used for an in-depth analysis of protein-protein interactions (PPI) through ZDOCK(78). ZDOCK v3.0.2 was utilized to conduct rigid-body docking, where PrpA was designated as the receptor and Tpl2 was employed as the ligand. A final conformational sampling was carried out with an angular step size of 6 Å. Subsequently, 2000 poses were generated for each docking process and were clustered using an RMSD cut-off value of 6 Å and an interface cut-off of 9 Å, with the maximum number of clusters set at 60. The poses were ranked based on ZDOCK score. Following this, the 2000 predicted docked conformations underwent refinement and re-ranking using the RDOCK module to eliminate clashes and optimize polar and charge interactions, aiming to identify a near-native conformation. The final- docked conformation was selected from each docking process based on having the best ZDOCK and RDOCK scores among all refined conformations.

### In Silico Mutagenesis of Conserved Interface Residues

To scrutinize the influence of specific residues on the binding affinity of a protein-protein complex, computational mutagenesis is widely applied, considering the intricate structure of the complex. This approach entails mutating residues situated at the interface of the wild-type (WT) complex structure, followed by estimating the binding affinity of the resulting complex(79, 80). As per the predictions made by the Build and Edit method, the residues Trp309, Glu103, and Glu129, participating in the interaction between PrpA and Tpl2, underwent mutation. Specifically, Arg, Trp, and Arg substitutions were introduced for these residues. It is anticipated that these mutations will result in a reduction in the affinity between PrpA and Tpl2.

### Molecular Dynamics Simulation and Binding Free Energy Calculation

To delve deeper into the molecular mechanisms of interaction between PrpA and Tpl2, Gromacs 2020 software was employed for molecular dynamics (MD) simulations of the PrpA-Tpl2 structural complex. The topology parameters for the structural complexes were generated using the AMBER99SB-ILDN forcefield(81). The TIP3P explicit water model was selected, with a minimum distance of 1.0 nm between protein atoms and the water box edge. Sodium or chloride ions were added to neutralize the system charge based on docking results. The molecular dynamics simulation workflow comprised four key steps: energy minimization, heating, equilibration, and production dynamics simulation. Initially, the protein heavy atoms were constrained, and energy minimization of water molecules was performed, utilizing 5000 steps each of the steepest descent and conjugate gradient methods over 10,000 steps. This process aimed to optimize the system’s energy and resolve any steric clashes. Subsequently, constraints were released, and the entire system underwent another 10,000-step energy minimization. For heating, the system was gradually heated to 300 K over 50 ps, followed by a 50 ps equilibration under the NPT ensemble. Finally, a 100 ns molecular dynamics simulation was conducted under the NPT ensemble, saving trajectory data every 10 ps for subsequent analysis using the trajectory module. The stability and behavior of the structural complexes were assessed by calculating the Root Mean Square Deviation (RMSD) and Root Mean Square Fluctuation (RMSF) over a 100 ns simulation run. These analyses were conducted to monitor the output of the Molecular Dynamics (MD) simulation and evaluate the overall structural stability and fluctuation of the complexes over the simulation time. Additionally, a comprehensive analysis of system dynamics was conducted through visualization using the Visual Molecular Dynamics (VMD) program(82) and DS. The free intermolecular binding energy of Tpl2 with PrpA or PrpA mutant was estimated using the Molecular Mechanics/Poisson Boltzmann Surface Area (MM/PBSA) method. This approach calculates the binding free energy by considering the molecular mechanics energy, solvation free energy, and the entropy contribution, providing valuable insights into the strength of the interactions between Tpl2 and PrpA or its mutant(83). The binding free energy (BFE) ΔGbind for the complex was calculated according to the below equation for the WT and mutated structural complexes(84).

Δ*Gbind* = *Gcomplex* – (*Gprotein1* + *Gprotein2*)

The *G_complex_* denotes the binding energy of the PrpA/Tpl2 structural complex, whereas *G_protein1_*and *G_protein2_* represent the energies of the constituent proteins within the complex. The estimation of the free energy of binding was conducted using the *g_mmpbsa* tool of GROMACS, employing the MM/PBSA method(85).

### Construction of Site-Mutagenesis Plasmid for *prpA*

Three mutations were introduced by altering residues W309R, E103W, and E129R through the PCR method. In brief, heat-killed *B. melitensis* M5-90 was used as a DNA template, and the primers for obtaining the mutated *prpA* product were as follows: *prpA*-mutation forward 5’- ATACGACTCACTATAGGGAGACCCAAGCTGGCTAGCATGGCAAGACATTCCTTCTTCTGCGTCGATGGGC-3’; *prpA*-mutation reverse 5’ TGATCAGCGGGTTTAAACGGGCCCTCTAGACTCGAGTTACTTGTCATCGTCGTCCTTGTAATCTGCCATG-3’. The reaction conditions were as follows: initial denaturation at 96 °C for 3 min followed by 23 cycles of 95 °C for 15 s, annealing at 58 °C for 15 s, and 72 °C for 20 s; the final extension was 72 °C for 1 min. The obtained PCR products were inserted into the pcDNA3.1- Flag-C, and the restricted enzyme sites were MluI and XhoI. The construct was verified by sequencing.

### Flow Cytometric Analysis to Evaluate CD4+ and CD8+ T Cells

Flow cytometric analysis of CD4+ and CD8+ T cells was performed in splenocytes from mice infected for 30 and 45 days with *B. melitensis* M5-90 or *B. melitensis* M5-90 *prpA* mutant. After passing the spleen cells through a 100-μm cell strainer and treating the samples with ACK buffer (0.15 M NH_4_Cl, 1.0 mM KHCO_3_, 0.1 mM Na_2_EDTA [pH 7.2]) to lyse red blood cells, splenocytes were washed with PBS. Following cell counting, 2 × 10^6^ cells/mouse were re-suspended in PBS (200 μL) and stained with a cocktail of anti-CD4 FITC (Biolegend, San Diego, CA) and anti-CD8a PE (Biolegend, San Diego, CA). The cells were incubated at 4 °C for 1 h and then washed with PBS. Flow cytometry analysis was performed using a BD FACSAria III (BD Biosciences, USA), and data were collected for 1.2 × 10^5^ cells/mouse. Data were analyzed using Flowjo software (Tree Star, Ashland, OR, USA).

### Mice Vaccination and Challenge

#### Clearance in Mice

Four to six-week-old Balb/c female mice were procured from the same institute mentioned previously. The animals were housed in a biosafety level 3 laboratory and provided ad libitum access to food and water. Mice were subcutaneously inoculated with 2 × 10^5^ CFU/0.2 mL of PBS diluent of *B. melitensis* M5-90 or *B. melitensis* M5-90 *prpA* mutant and *B. melitensis* M5-90 *prpA* complement strain (*B. melitensis* M5-90 prpA-c) or 0.2 mL PBS (uninfected vehicle control). At designated time intervals post-infection, mice were humanely euthanized following institutional animal care protocols. To quantify the CFU of *Brucella*, individual spleens were isolated, macerated in 1 mL of PBS, serially diluted, and then plated in duplicate on *Brucella* selective agar (containing 5% equine serum from HyClone, USA, and Farrell antimicrobials from OXOID^TM^, UK). After an incubation period of 5 – 7 days at 37°C, the CFU count was determined. **Protection in Mice:** To assess the ability of strain *B. melitensis* M5-90 *prpA* mutant to confer protection against a *B. melitensis* M28 challenge, female Balb/c mice (n = 6) were vaccinated with 2 × 10^5^ CFU/0.2 mL of PBS diluent of *B. melitensis* M5-90 or *B. melitensis* M5-90 *prpA* mutant and 0.2 mL PBS (uninfected vehicle control). Seven weeks post-vaccination, mice were intraperitoneally (i.p.) challenged with 2 × 10^5^ CFU wild-type *B. melitensis* M28. Two weeks post-challenge, mice were euthanized, and splenic bacterial CFU were determined.

### Histological Analysis

Spleen and liver tissues were isolated and fixed in 4% formalin. Five μm thick tissue specimens were meticulously prepared for subsequent mounting onto glass slides. Following established protocols, the specimens underwent Hematoxylin and Eosin staining, with microphotography facilitated using an Olympus microscope (BX53F model, Japan) and digital imaging software (analysis TS Olympus Corp., Japan). Subsequently, the enumeration of hepatic microgranuloma foci was conducted across 10 distinct microscopic fields (400× magnification).

### Statistical Analysis

The data are presented as mean ± standard error of the mean (SEM) (standard deviation [SD]), and all experiments were repeated at least three times. Statistical analyses were conducted using R (v4.0.5) with the “tidyverse” package and unpaired Student t-test. Significance levels are indicated in the figures as follows: (* P < 0.05, ** P < 0.01, *** P < 0.005, and ns = not significant).

## Competing interest

The authors declare no competing interest.

## Author contributions

Conceptualization: Chuangfu Chen, Jihai Yi and Zhongchen Ma.

Writing – review & editing: Huan Zhang, Yong Wang, Yuanzhi Wang, Jinliang Sheng, Hui Zhang and Kait Zhumanov.

Data curation: Huan Zhang Xujin Xia, Mingguo Xu and Zhen Wang

Resources: Jihai Yi, Zhongchen Ma and Chuangfu Chen

Investigation: Huan Zhang, Yueli Wang, Huilin Hou, Yaqian Wang, Xiaoyu Deng, Zhenyu Xu, Zhongchen Ma and Changsuo Zhang.

Writing – Original Draft preparation: Huan Zhang, Yuanzhi Wang. Software: Huan Zhang.

## Funding

This work was supported by the National Natural Science Foundation of China (to Chuangfu Chen, grant no. U1803236), the Youth Innovation and Talent Training Program of Shihezi University (to Jihai Yi, grant no. CXPY202109), and the Shihezi University International Science and Technology Cooperation Promotion Program (to Jihai Yi, grant no. GJHZ202203).

## Acknowledgments

We thank MedSci (www.medsci.cn) for its linguistic assistance during the preparation of this manuscript.

## Notes

### Competing Interest Statement

The authors have declared no competing interest.

